# Temporal association activates projections from the perirhinal cortex and ventral CA1 to the prelimbic cortex and from the prelimbic cortex to the basolateral amygdala

**DOI:** 10.1101/2023.08.16.553604

**Authors:** Thays Brenner dos Santos, Juliana Carlota Kramer-Soares, Cesar Augusto de Oliveira Coelho, Maria Gabriela Menezes Oliveira

**Author notes:** Corresponding author: Thays Brenner dos Santos, Ph.D. **Author contributions:** TBS conceptualized the study and methodology, performed the investigation, conducted the data curation, formally analyzed and visualized the data, administrated the project, and wrote the original draft. JCK, CAOC, and MGMO reviewed the draft and supervised the study. TBS, JCK, CAOC, and MGMO acquired funding.

## Abstract

In temporal associations, the prelimbic cortex (PL) has persistent activity during the interval between the conditioned stimulus (CS) and the unconditioned stimulus (US), which maintains a CS representation. Regions cooperating for this function or encoding the CS before the interval could neuroanatomically connect to the PL, supporting learning. The basolateral amygdala (BLA) has CS- and US-responsive neurons, convergently activated. The PL could directly project to the BLA to associate the transient CS memory with the US. We investigated the neural circuit supporting temporal associations using the CFC-5s task, in which a 5-second interval separates the contextual CS from the US. Injecting retrobeads, we quantified c-Fos in PL- or BLA-projecting neurons from 9 regions after CFC-5s or contextual fear conditioning (CFC), in which CS/US overlap. The CFC-5s activated ventral CA1 (vCA1) and perirhinal cortex (PER) neurons projecting to the PL, and PL neurons projecting to BLA. Both CFC-5s and CFC activated vCA1 and lateral entorhinal (LEC) neurons projecting to BLA, and BLA neurons projecting to PL. Both conditioning activated the PER, LEC, cingulate and infralimbic cortices, nucleus reuniens, and ventral subiculum. Results added new relevance to the PER→PL projection and showed that the PL/BLA are reciprocally functionally connected in CFC-5s.

## 1. Introduction

In associations of stimuli separated in time, a neural representation of the conditioned stimulus (CS) is maintained over a time interval to be associated with a later unconditioned stimulus (US). They are distinct from associations of stimuli overlapped in time, in which the CS and US coincide in time, not requiring a CS representation because there is no time interval between the stimuli, and the CS is not absent from the environment during the US delivery (Pavlov 1927; Raybuck and Lattal 2014).

The prelimbic cortex (PL), a subdivision of the medial prefrontal cortex (mPFC), is required for temporal associations, such as trace fear conditioning and contextual fear conditioning with a 5-second interval (CFC-5s), in which an auditory or contextual CS are separated in time from the US, respectively (Gilmartin and Helmstetter 2010; Guimarãis et al. 2011; Gilmartin et al. 2012; Santos et al. 2017; Mukherjee and Caroni 2018; Rose et al. 2021). In contrast, the PL is not required for learning associations overlapped in time, such as auditory (AFC) or contextual fear conditioning (CFC, Corcoran and Quirk 2007; Gilmartin and Helmstetter 2010; Guimarãis et al. 2011; Santos et al. 2017), but PL encoding ensembles would reorganize over time to support remote fear memories (DeNardo et al. 2019). Persistent activity is observed in the PL during the time interval in trace fear conditioning (Baeg et al. 2001; Gilmartin and McEchron 2005) and PL inhibition during the time interval impaired learning (Gilmartin et al. 2013), suggesting that the PL has a role related to the time interval, which could be supporting a transient CS memory to be used in the future. This interpretation is also favored by working memory (WM) studies, in which persistent activity in the mPFC during the delay period between the sample and choice phases, that is, between a cue and a goal-directed response, is also observed (Liu et al. 2014; Kamigaki and Dan 2017; Abbas et al. 2018).

In turn, the basolateral amygdala (BLA) has neurons responsive to both the CS and US that are convergently activated during fear conditioning, supporting their association (Barot et al. 2009; Wolff et al. 2014; Gore et al. 2015). The BLA is required for associations overlapped or nonoverlapped in time (Guimarãis et al. 2011; Gilmartin et al. 2012; Kwapis et al. 2011; Chau et al. 2013; Kochli et al. 2015; Kirry et al. 2020; Sun et al. 2020). In temporal associations, brain regions encoding the CS could functionally and neuroanatomically connect to brain regions supporting its maintenance over time by persistent activity, such as the PL. On the other hand, brain regions supporting the CS representation over time could functionally and neuroanatomically connect to brain regions associating the CS with the US, such as the BLA. Activation of projections to the PL could converge the CS to be maintained during the time interval, and activation of projections to the BLA could convey the CS representation to be associated with the US after the time interval. In line, the PL has direct connections to the anterior subdivision of the BLA through glutamatergic ipsilateral and contralateral projections, making synapses onto pyramidal neurons and interneurons (Rosenkranz et al. 2001; Hübner et al. 2014). In turn, the posterior subdivision of the BLA projects back to the PL (Cheriyan et al. 2016).

We investigated the activation of direct projections to the PL or BLA supporting temporal learning using the CFC-5s task. To observe projection-specific activation, we used the retrograde tracer retrobeads (RB) and quantified learning-related c-Fos expression in neurons projecting to the PL (Experiment 2) or the BLA (Experiment 3) from the PL (Experiment 3), BLA (Experiment 2), the cingulate cortex (AC); infralimbic cortex (IL); ventral CA1 (vCA1); ventral subiculum (vSUB); lateral entorhinal cortex (LEC); perirhinal cortex (PER), nucleus reuniens of the thalamus (RE) and ventral striatum (VS, Experiments 2 and 3) following the CFC-5s or CFC training. We hypothesized that a direct connection from the PL to the BLA would contribute to CFC-5s learning and direct connections from brain regions related to forming a contextual representation to the PL.

The vCA1 and vSUB send glutamatergic ipsilateral projections to the PL (Jay et al. 1992; O’Mara et al. 2006; Hoover and Vertes 2007), in contrast to the dorsal hippocampus (DH), which have undetected to sparse direct projections (Hoover and Vertes 2007; Kim and Cho 2017; Ye et al. 2017). The PER also projects to the PL (Hwang et al. 2018). The ventral hippocampus (VH) is required for contextual learning (Rudy and Matus-Amat 2005), and both the VH and PER for CFC (Bucci et al. 2000; Bast et al. 2001; Zhang et al. 2001) and trace fear conditioning (Rogers et al. 2006; Yoon and Otto 2007; Kholodar-Smith et al. 2008b; Czerniawski et al. 2011; Czerniawski et al. 2012; Suter et al. 2013). Due to its facilitated connection compared to the DH, the vCA1, vSUB, or PER could activate projections to the PL to convey a contextual CS representation.

The LEC and RE can also mediate hippocampal-cortical interactions. The RE has single or collateral projections to CA1 and PL, which reciprocally project back to the RE (Hoover and Vertes 2012; Varela et al. 2014). The LEC receives projections from the CA1 (Insausti et al. 1997) and projects to the PL (Insausti et al. 1997; Delatour and Witter 2002). Both the EC and the RE are required for trace fear conditioning (Esclassan et al. 2009; Eleore et al. 2011; Lin et al. 2020; East et al. 2021; Wu and Chang 2021; Jhuang and Chang 2023), and the RE for functional interaction between the DH and PL in spatial navigation (Ito et al. 2015) and spatial WM (SWM, Hallock et al. 2016). In addition, the PER (Navaroli et al. 2012) and EC (Egorov et al. 2002) have neurons that change their firing in response to a stimulus, endogenously maintained stable after the stimulus withdrawal. This feature could also support CS transient memories in cooperation with the PL.

The AC and IL are reciprocally connected to the PL (Hoover and Vertes 2007) and have CS- or interval-responsive neurons during trace fear conditioning but in a different activity pattern from the PL (Baeg et al. 2001; Gilmartin and McEchron 2005; Steenland et al. 2012; Song et al. 2015), and could constitute a mPFC internal circuit to support CS transient memories. The vCA1, vSUB, PER, RE, LEC, AC, and IL also project to the BLA (Canteras and Swanson 1992; Pitkänen et al. 2000; Hintiryan et al. 2021). We also quantified c-Fos- and RB-positive cells in the ventral striatum (VS) as a control for unspecific diffusion, given that it does not project to the PL or the BLA (Haber 2011).

PL→BLA and vCA1→PL connections have been studied in trace fear conditioning. Pre-training chemogenetic or optogenetic inhibition during the time interval of PL inputs to the BLA did not impair trace fear conditioning. Only pre-training chemogenetic inhibition of the PL→BLA projection using weak training (2 CS-US pairings) decreased the CR compared to a non-inhibited group (Kirry et al. 2020). The CFC-5s training, encoded with 1 CS-US pairing and in which the CS is already the context, not having other predictive stimuli, such as the context in trace fear conditioning (Santos et al. 2017), may be a more suitable task to investigate this connection. In addition, comparing an association overlapped in time (CFC group) may better indicate if this connection is related to temporal learning. In turn, the opto-inhibition of vCA1 inputs to the PL during training also did not impair trace fear conditioning (Twining et al. 2020), suggesting that PL afferences may be related to the CS modality. If CFC-5s, which used a hippocampus-dependent contextual CS, engages vCA1→PL connection in opposition to trace fear conditioning, it strengthens the idea that this projection is functional to convey the CS to the PL in temporal associations.

### 2. Materials and Methods

#### 2.1 Subjects

Experiments were performed on thirty-six male *Wistar* rats 10 weeks old at the start of the Experiments, weighing from 250 to 330 g. Rats were housed in groups of 4 and obtained from CEDEME (Center of Development of Experimental Models for Medicine and Biology, Universidade Federal de São Paulo, Brazil). All animals were acclimatized for one week and housed in transparent polysulfone plastic cages (44 × 35 × 20 cm) individually ventilated, with corn-cob bedding on the floor. Room temperature was controlled (22 °C ± 1 °C), and a light-dark cycle was maintained (lights on at 07:00 am and off at 07:00 pm). Food and water were available *ad libitum*. We assessed the rat’s welfare daily. The Ethical Committee for Animal Research of Universidade Federal de São Paulo approved the study (number #6790140616). All procedures followed the policies and guidelines of the National Council for the Control of Animal Experimentation (CONCEA, Brazil) and the NIH Guide for the Care and Use of Laboratory Animals. The sample size was estimated on G*Power (Faul et al. 2007) to ensure adequate power to detect effect sizes previously observed using the CFC-5s training (Santos et al. 2023).

#### 2.2 Retrograde tracer

For the retrograde labeling of activated neurons projecting to the PL or the BLA, we used fluorophore coated latex microspheres referred to as red (excitation = 540-552 nm, emission = 590 nm) retrobeads (RBs, R180-100, Lumafluor, Naples, US). We chose RBs due to their limited diffusion, high sensitivity of labeling cells and low dendritic labeling, excellent fading resistance after infusion, exclusive retrograde travel, and minimal entry to undamaged fibers of passage (Schofield 2008). The volumes used were based on previous studies on the PL and BLA (Song et al. 2015; Marek et al. 2018).

#### 2.3 Stereotaxic surgery

Rats underwent surgery to inject the RBs in the PL or BLA. Rats were anesthetized intraperitoneally (IP) with Fentanyl (0.02 mg/kg, Cristália, Sao Paulo, BR), Ketamine (90 mg/kg, Syntec, Santana do Parnaíba, BR), and Xylazine (10 mg/kg, Syntec) and then fixed in the stereotaxic frame (David Koft Instruments, Tujunga, US) by the upper incisors and ear bars. We shaved their heads, covered their eyes with ophthalmic gel, and maintained the body temperature at 37°C with a heating pad. After asepsis and subcutaneous administration of 3% lidocaine hydrochloride in conjunction with norepinephrine (Syntec), the scalp was exposed, the periosteum retracted, and the bregma and lambda were placed in the same horizontal plane. We used a 2.0 μl syringe (Hamilton, Reno, US) to bilaterally infuse a volume of 0.4 μl of the RBs in the PL (anteroposterior, AP, +3.2 mm, ML ±0.7 mm, DV -3.5 mm, from bregma) or of 0.2 μl of the RBs in the BLA (AP -3.0 mm, ML ±5.0 mm, DV -8.3 mm, from bregma), according to Paxinos and Watson (2007). The speed of the infusion was 0.1 μl/min. The needle remained in the target regions for 10 min to prevent back-flow. The scalp was sutured, and a topical antibiotic was applied to the surgical area. All rats received a subcutaneous injection of 0.1 ml 0.9% Saline and intramuscular injections of 0.1 ml Pentabiotic (Zoetis, Parsippany-Troy Hills, US) and 0.1 ml Meloxicam (Ourofino, Cravinhos, BR), which was also administered the following two days. Animals recovered from surgery for at least one week.

#### 2.4 Apparatus

The conditioning chamber consisted of a 22 × 27 × 45 cm box with black acrylic walls, a clear acrylic top, and a video camera attached to the top (AVS Projetos, São Paulo, BR). The shock grid comprised parallel stainless-steel bars of 0.4 cm diameter spaced 1.2 cm apart and connected to an electric generator. We cleaned the conditioning chamber with a 20% ethanol solution after the training session for each animal. The transport cage consisted of a 22 × 35 × 20 cm cage with corn-cob bedding from the homecage of the rat and a clear polysulfone cover topped with a flannel. It transported the animals from the homecage to the experimental room, and it was the context during the 5-second interval in the CFC-5s training.

#### 2.5 Behavioral procedures

## 2.5.1 Habituation

Habituations were conducted in the experimental room for three consecutive days before the training sessions. We handled each rat for 5 min in two transport cages, moving them from one to another every 15 s to familiarize the animals with the experimental conditions and experimenter.

## 2.5.2 Contextual fear conditioning with a 5-second interval (CFC-5s)

CFC-5s was conducted as previously described (Santos et al. 2017; Santos et al. 2020; Santos et al. 2023). Rats were pre-exposed to the conditioning chamber for 5 min, placed in the transport cage beside the conditioning chamber for a 5-second interval, and then re-exposed to the conditioning chamber, delivering one immediate footshock (0.8 mA, 1 s). We returned them immediately to their homecage in the transport cage.

## 2.5.3 Contextual fear conditioning (CFC)

CFC was performed as previously described (Santos et al. 2017; Santos et al. 2020; Santos et al. 2023). We placed the rats in the conditioning chamber for 5 min delivering one footshock (0.8 mA, 1 s) at the end. We removed the animals immediately and returned them to their homecage in the transport cage.

### 2.6 c-Fos immunofluorescence

Ninety minutes after the training sessions, we anesthetized the rats with IP injections of Lidocaine (10 mg/kg; Bravet, Rio de Janeiro, Brazil) and Thiopental (150 mg/kg; Cristália). Next, we transcardially perfused them with 0.9% saline at 4 °C for 1 min, followed by 4% PFA (paraformaldehyde; Synth, Diadema, Brazil) at 4 °C for 15 min at 12 ml/min. A peristaltic pump (Cole Parmer, Vernon Hills, US) drove the perfusions. We injected 0.1 ml of heparin (Cristália) directly into the left ventricle. We removed the brains from the skull, post-fixed them in 4% PFA for 24 h at 4 °C and transferred them to 30% sucrose (Synth) in 0.02 M KPBS (potassium phosphate-buffered saline) solution until the samples sank (48 h). We freeze the brains with dry ice and store them at -80 °C. We obtained coronal sections of 30 μm in five series using a cryostat at -20 °C (Leica CM1850, Wetzlar, Germany). We stored the series in an anti-freezing solution with ethylene glycol (Synth) and 30% sucrose at -20 °C.

First, free-floating sections of one series were washed in 0.02 M KPBS 3 times for 10 min each. Next, sections were transferred to a solution containing 1% hydrogen peroxide for 15 min at room temperature, rewashed, and transferred to a blocking solution containing 2% normal goat serum (NGS, S-1000-20, Vector Laboratories, Burlingame, US) for 1 h at room temperature. Then, sections were incubated in the primary antibody solution containing a rabbit anti-c-Fos antibody (1:4000; ab190289, Abcam, Cambridge, UK), 2% NGS, and 0.3% Triton X-100 (85111, ThermoFisher) for 48 h at 4 °C. Sections were washed and transferred to a secondary antibody solution containing goat anti-rabbit Alexa Fluor 488 (1:500, A11008, ThermoFisher) and 0.3% Triton X-100 for 2 h at room temperature. Finally, we washed the sections, counterstained them with 4’6’-diamidino-2-phenylindole (DAPI) for 10 min (1:10000, D9542, Sigma-Aldrich), and mounted them on gel-coated slides coverslipped with Vectashield fluorescent mounting medium (H-2000, Vector Laboratories). We selected at least one sample of each group for the batches of immunofluorescence reactions and performed a reaction without the primary antibody as a negative control.

### 2.7 Image analysis

We imaged sections on a fluorescent microscope (Olympus, BX50, Waltham, US) at 20x magnification and fixed image size (0.87 mm × 0.66 mm) and area (0.5742 mm^2^). Smaller brain regions were delimited using the ImageJ software (NIH, Washington, United States), excluding adjacent areas from the cell counting. More extensive brain regions were obtained in more than one frame. We took reference images at 4x magnification for assistance. Cell counting was standardized for cells/0.5742 mm^2^. An experimenter, blind to the experimental group of the animals, captured 6 images from each brain region from 3 AP coordinates (one anterior, one intermediate, and one posterior) of both hemispheres. We took 6 pictures of the PL, AC, and IL (+3.72, +3.00, +2.76 mm AP from bregma); RE (-1.80, - 1.92, -2.04 mm AP from bregma); BLA and VS (-2.28, -2.64, -3.00 mm AP from bregma); PER, vCA1, and vSUB (-5.40, -5.64, -5.88 mm AP from bregma) and LEC (-6.96, -7.20, - 7.44 mm AP from bregma).

### 2.8 Cell quantification and Injection site verification

We counted the number of DAPI-positive, c-Fos-positive, RB-positive, and double-labeled cells (colocalization of RB-positive and c-Fos-positive cells) in ImageJ software using the package Fiji, blinded to the experimental groups. Double-labeled cells resulted in the center and surrounded labeled with c-Fos and RB, respectively. Counts were averaged per animal. Serial brain sections were analyzed to determine if the infusion of the retrograde tracer was localized in the PL and the BLA. Only animals with the infusions restricted to the target brain regions were included in the statistical analysis and reported. We adapted coronal sections (Paxinos and Watson 2007) and drew images to illustrate the injection sites in the Photoshop CS6 software (Adobe, San Jose, US).

### 2.9 Experimental design

In Experiment 1 (n = 30, 6/group) rats were habituated for 3 consecutive days and the next day training in the CFC-5s or CFC, together with 3 control groups. In the training session, the US group was exposed to the footshock (0.8 mA, 1s) immediately in the conditioning chamber, controlling for US perception and non-associative learning; the CT group was exposed to the conditioning chamber for 5 min, without receiving any footshock, controlling for contextual learning and exploration, and the CT-5s group was exposed to the conditioning chamber for 5 min, removed for a 5-second interval, and then returned to the conditioning chamber, without receiving any footshock, controlling for the time interval and the manipulation that it requires. Forty-eight hours later, all groups were tested in the conditioning chamber and freezing time was measured manually with a chronometer to infer conditioned responses (CR). In Experiments 2 and 3, rats received bilateral injections of RBs into the PL (Experiment 1, n = 18, 6/group) or the BLA (Experiment 2, n = 18, 6/group). After recovering for one week, they were habituated for 3 consecutive days and the next day training in the CFC-5s or CFC. All animals were euthanized 90 min later for c-Fos immunofluorescence, including a homecage (HC) group that stayed in the homecage during the training session. All behavioral procedures were conducted from 10:00 am to 2:00 pm, and groups were distributed evenly throughout the period. The order of removal from their homecage (first to fourth) was also evenly distributed among the groups. We ensured that each homecage had animals from different groups.

### 2.10 Statistical analysis

Data were analyzed by Generalized Linear Models (GZLM), a generalization of General Linear Models used to fit regression models for univariate data presumed to follow the exponential class of distributions (McCullagh and Nelder 1989). Estimations were adjusted to Linear, Gamma, or Tweedie probability distributions with identity link functions according to the Akaike Information Criterion (AIC). GZLM evaluated the main effect of the group in the freezing time in the training session, DAPI-positive cells, c-Fos-positive cells (standardized by DAPI-positive cells), RB-positive cells (standardized by DAPI-positive cells), and double-labeled cells (standardized by RB-positive cells) in rats injected with RBs in the PL (Experiment 2) or the BLA (Experiment 3), in each one of the nine brain regions investigated. Estimations were considered statistically significant if p < 0.050. In these cases, we used the LSD test when necessary (IBM SPSS Statistics, 23). We also compared effect sizes using standardized regression coefficients (β). Values above 0.35 are considered large (Cohen 1992). P-values around 0.05 in the post hoc comparisons with large effect sizes are interpreted as a trend toward statistical significance. We created all graphs in GraphPad Prism 8 (GraphPad, San Diego, US).

#### 3. Results

### 3.1 CFC and CFC-5s learning elicited conditioned responses

Both CFC and CFC-5s training induced higher freezing time in the conditioned context than the context and US exposures alone, suggesting learning of the contextual fear association (Experiment 1, Figure 1). GZLM showed a significant effect of the group in the freezing time during the test (W = 114.520; df = 4; p = 0.001), but not in the training session (W = 2.023; df = 3, p = 0.568). LSD test showed that CFC and CFC-5s groups had a higher freezing time than the US (CFC p = 0.001; β = 1.172; CFC-5s p = 0.001; β = 1.196), CT (CFC p = 0.001; β = 1.833; CFC-5s p = 0.001; β = 1.857), and CT-5s (CFC p = 0.001; β = 2.029; CFC-5s p = 0.001; β = 2.052) groups. The US group also had a higher freezing time than the CT (p = 0.011; β = 0.661) and CT-5s (p = 0.001; β = 0.857) groups.

**Figure 1.**
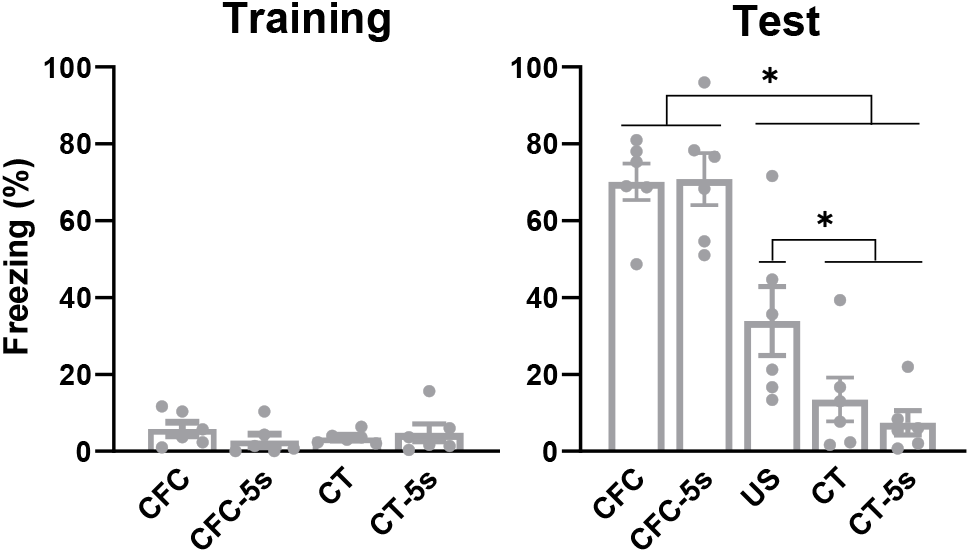
CFC and CFC-5s training produced conditioned responses. Mean (± standard error) of freezing time in the training **(A)** and test **(B)** sessions in CFC, CFC-5s, US, CT, and CT-5s groups. * Indicates p < 0.050. Generalized Linear Models followed by LSD test. Dots show sample distribution. CFC: contextual fear conditioning; CFC-5s: CFC with a 5-second interval; US: unconditioned stimulus; CT: context.

### 3.2 CFC-5s learning induced the activation of PER and vCA1 projections to the PL

There was a significantly higher c-Fos expression in PL-projecting neurons in the vCA1 and the PER induced by CFC-5s training compared to CFC and HC groups and significantly higher c-Fos expression in PL-projecting neurons in the BLA and LEC induced by both conditioning compared to HC group (Experiment 2, Figures 2D-G). GZLM showed an effect of the group for the double-labeled neurons in the BLA (W = 11.107; df = 2; p = 0.004), vCA1 (W = 8.026; df = 2; p = 0.018), PER (Wald = 13.193; degree of freedom = 2; p = 0.001), and LEC (W = 9.270; df = 2; p = 0.010), but not in the AC (W = 5.026; df = 2; p = 0.081), IL (W = 2.286; df = 2; p = 0.319), RE (W = 2.407; df = 2; p = 0.300), vSUB (W = 733; df = 2; p = 0.094), and VS (mean ± standard error = 0.000 ± 0.000 in all groups). LSD test showed that the CFC-5s group had a significantly higher value of double-labeled neurons compared to CFC and HC groups in the PER (CFC: p = 0.013; β = 1.063; HC: p = 0.001; β = 1.506), to the HC group in the vCA1 (p = 0.005; β = 1.297), and a trend toward statistical significance, with large effect size, to the CFC group in the vCA1 (p = 0.062; β = 0.871). The CFC and CFC-5s groups had a significantly higher value of double-labeled neurons than the HC group in the BLA (CFC p = 0.003; β = 1.293; CFC-5s p = 0.005 β = 1.253). The CFC group also had a significantly higher value of double-labeled neurons than the HC group in the LEC (0.003; β = 1.375), and the CFC-5s group had a trend toward statistical significance, with a large effect size than the HC group in the LEC (p = 0.061; β = 0.853).

**Figure 2.**
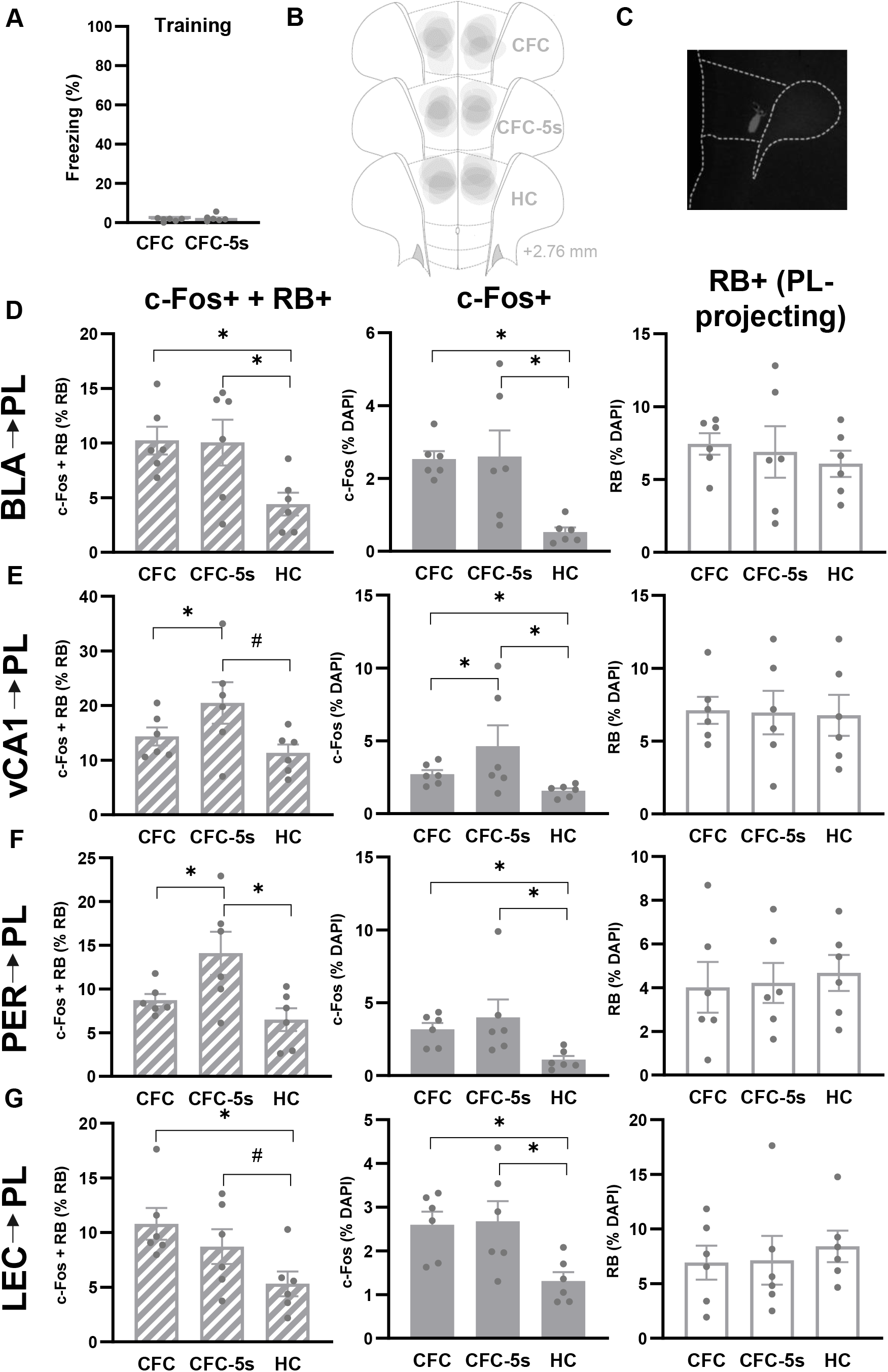
CFC-5s activated vCA1 and PER projections to the PL. **(A)** Mean (± standard error) of freezing time in the CFC and CFC-5s groups training session. **(B)** Retrobeads (RBs) diffusion in the PL in the CFC (contextual fear conditioning), CFC-5s (CFC with 5-second interval), and HC (homecage) groups. **(C)** Representative image of RBs in the PL. Mean (± standard error) of double-labeled (c-Fos-positive and RB-positive) cells (left, striped), c-Fos-positive cells (middle, grey), and RB-positive cells (right, white) in the **(D)** basolateral amygdala (BLA), **(E)** ventral CA1 (vCA1), **(F)** perirhinal cortex (PER), and **(G)** lateral entorhinal cortex (LEC). * Indicates p < 0.050. # Indicates trend towards statistical significance; p = 0.062; β = 0.871 in **(E)** and p = 0.061; β = 0.853 in **(G)**. Generalized Linear Models followed by LSD test. Dots show sample distribution.

### 3.3 CFC-5s learning induced higher activation of the vCA1

There was a significantly higher c-Fos expression in the vCA1 induced by CFC-5s training compared to CFC and HC groups and induced by CFC training compared to the HC group. Both fear conditioning had a significantly higher c-Fos expression in the BLA, PER, LEC, AC, IL, RE, and vSUB than in the HC group (Figures 2D-G and 3A-D). GZLM showed an effect of the group for c-Fos-positive cells in the BLA (W = 40.005; df = 2; p = 0.001), vCA1 (W = 17.929; df = 2; p = 0.001), PER (W = 23.214; df = 2; p = 0.001), LEC (W = 16.563; df = 2; p = 0.001), AC (W = 21.259; df = 2; p = 0.001), IL (W = 29.112; df = 2; p = 0.001), RE (W = 24.289; df = 2; p = 0.001), and vSUB (W = 45.805; df = 2; p = 0.001), but not in the VS (W = 1.236; df = 2; p = 0.539). LSD test showed that the CFC-5s group had significantly higher c-Fos expression in the vCA1 than the CFC group (p = 0.048; β = 0.813). Both CFC and CFC-5s groups had significantly higher c-Fos expression than the HC group in the BLA (CFC p = 0.001; β = 1.461; CFC-5s p = 0.001; β = 1.411), vCA1 (CFC p = 0.044; β = 0.483; CFC-5s p = 0.001; β = 1.296), PER (CFC p = 0.002; β = 0.956; CFC-5s p = 0.001; β = 1.336), LEC (CFC p = 0.001; β = 1.260; CFC-5s p = 0.001; β = 1.340), AC (CFC p = 0.001; β = 1.623; CFC-5s p = 0.001; β = 1.192), IL (CFC p = 0.001; β = 1.517; CFC-5s p = 0.001; β = 1.678), RE (CFC p = 0.002; β = 1.113; CFC-5s p = 0.001; β = 1.476), and vSUB (CFC p = 0.001; β = 0.868; CFC-5s p = 0.001; β = 1.476).

**Figure 3.**
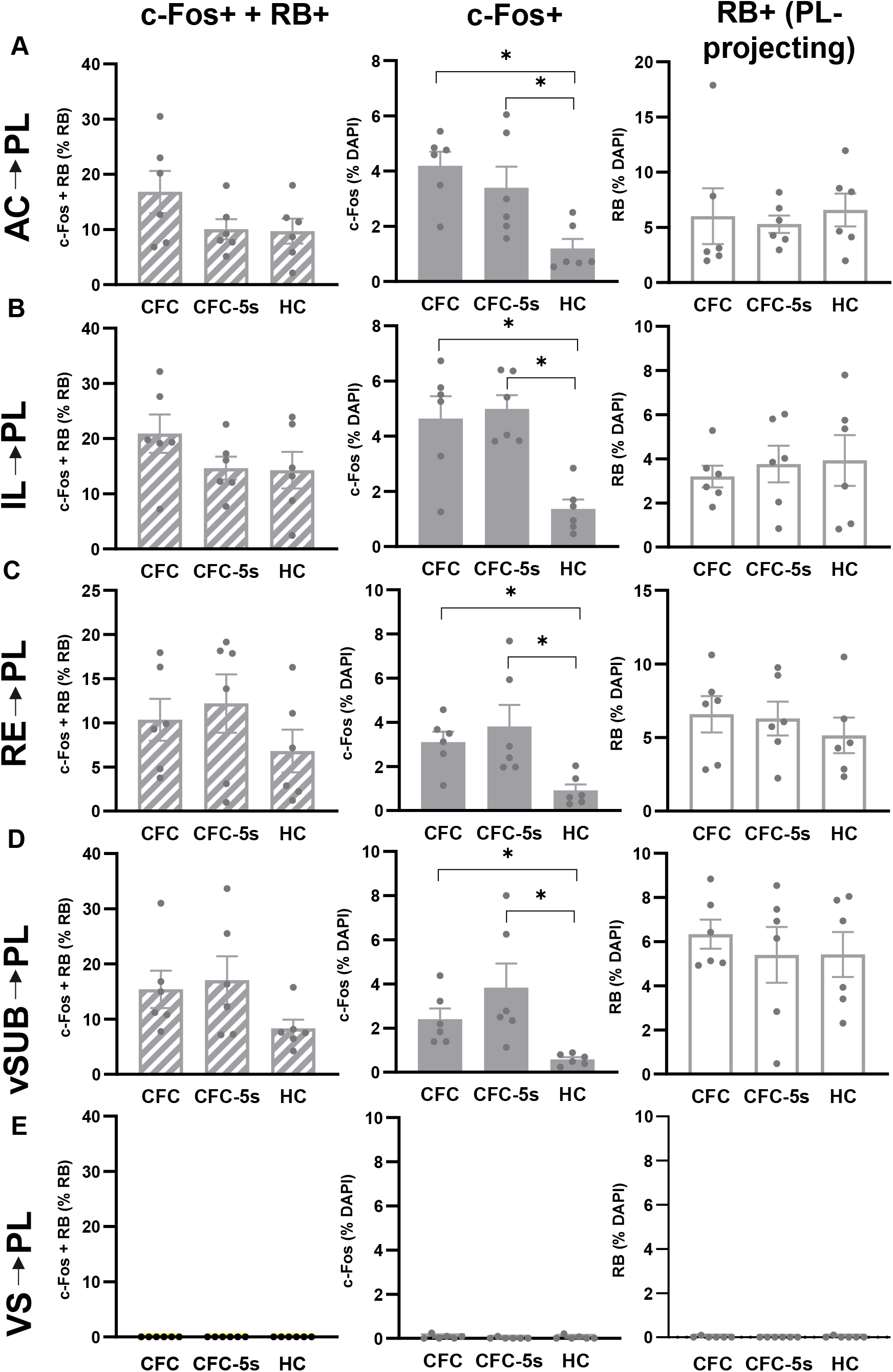
CFC and CFC-5s learning activated cells in the AC, IL, RE, and vSUB. Mean (± standard error) of double-labeled (c-Fos-positive and RB-positive) cells (left, striped), c-Fos-positive cells (middle, grey), and RB-positive cells (right, white) in the **(A)** cingulate cortex (AC), **(B)** infralimbic cortex (IL), **(C)** nucleus reuniens of the thalamus (RE), **(D)** ventral subiculum (vSUB), and **(E)** ventral striatum (VS). * Indicates p < 0.050. Generalized Linear Models followed by LSD test. Dots show sample distribution.

All groups had similar freezing times during the training session (Figure 2A) and an equal average of cells (DAPI-positive cells) and PL-projecting cells in all brain regions. GZLM did not show an effect of the group in the mean freezing time in the training session (W = 1.297; df = 1; p = 0.255) nor in DAPI-positive cells in the BLA (W = 2.693; df = 2; p = 0.260), vCA1 (W = 1.078; df = 2; p = 0.583), PER (W = 0.374; df = 2; p = 0.829), LEC (W = 0.629; df = 2; p = 0.730), AC (W = 2.283; df = 2; p = 0.319), IL (W = 0.107; df = 2; p = 0.948), RE (W = 0.824; df = 2; p = 0.662), vSUB (W = 3.990; df = 2; p = 0.136), and VS (W = 0.357; df = 2; p = 0.357). GZLM also did not show an effect of the group in RB-positive cells in the BLA (W = 0.711; df = 2; p = 0.701), vCA1 (W = 0.040; df = 2; p = 0.980), PER (W = 0.273; df = 2; p = 0.872), LEC (W = 0.474; df = 2; p = 0.789), AC (W = 0.416; df = 2; p = 0.812), IL (W = 0.457; df = 2; p = 0.796), RE (W = 0.937; df = 2; p = 0.626), vSUB (W = 0.678; df = 2; p = 0.712), and VS (W = 1.208; df = 2; p = 0.547). Overall, these results indicate that activation of vCA1 and PER projections to the PL and increased activation of vCA1 underlie CFC-5s learning, whereas activation of BLA projections to the PL, and increased activation of the BLA, PER, LEC, AC, IL, RE, and vSUB underlie CFC learning, despite of the time interval. Differential labeling of PL-projecting neurons among the groups could not explain the results. In addition, PL-projecting neurons in the VS were also undetected, suggesting specific PL targeting and labeling of PL-projecting neurons. Figures 2B and 2C illustrate the diffusion pattern of the RBs in the PL. Grey areas represented the locations of the micro-infusions.

### 3.4 CFC-5s learning induced the activation of PL projections to the BLA

There was a significantly higher c-Fos expression in BLA-projecting neurons in the PL induced by CFC-5s compared to CFC and HC groups and significantly higher c-Fos expression in BLA-projecting neurons in the vCA1 induced by both conditioning compared to the HC group (Experiment 3, Figures 4D and 4E). GZLM showed an effect of the group for the double-labeled neurons for the PL (W = 9.165 df = 2; p = 0.010) and vCA1 (W = 9.287 df = 2; p = 0.010) but not for the PER (W = 1.894; df = 2; p = 0.388), LEC (W = 0.006; df = 2; p = 0.997), AC (W = 2.707; df = 2; p = 0.258), IL (W = 0.142; df = 2; p = 0.931), RE (W = 2.810; df = 2; p = 0.245), vSUB (W = 4.572; df = 2; p =0.102), and VS (0.000 ± 0.000 for all groups). The LSD test showed that the CFC-5s group had a significantly higher value of double-labeled neurons compared to CFC and HC groups in the PL (CFC: p = 0.013; β = 1.130; HC: p = 0.006; β = 1.255). Both CFC and CFC-5s groups had a significantly higher value of double-labeled neurons than the HC group in the vCA1 (CFC: p = 0.015; β = 1.159; CFC-5s: p = 0.009; β = 1.281).

**Figure 4.**
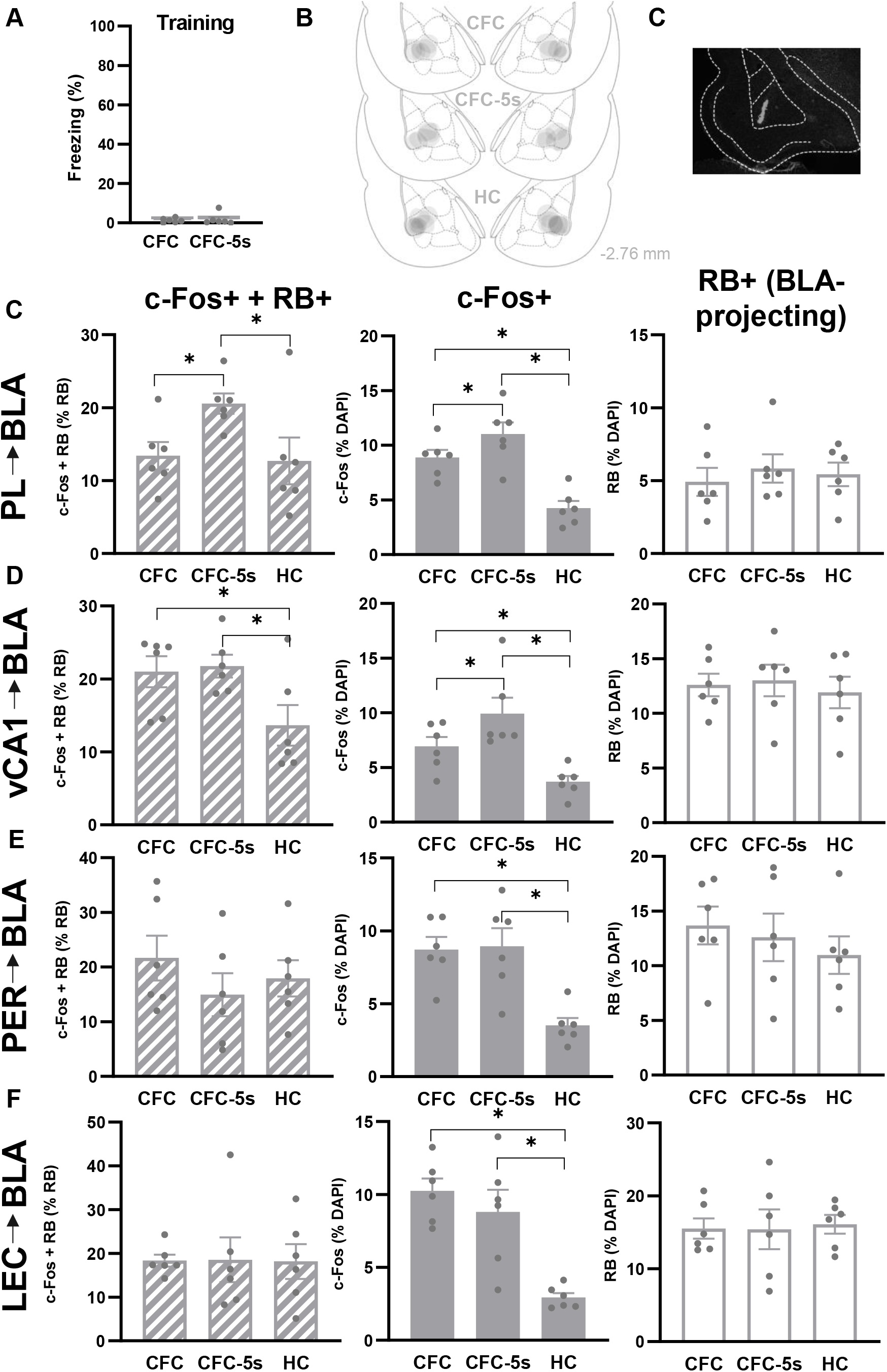
CFC-5s learning activated PL projections to the BLA. **(A)** Mean (± standard error) of freezing time in the CFC and CFC-5s groups training session. **(B)** Retrobeads (RBs) diffusion in the BLA in the CFC (contextual fear conditioning), CFC-5s (CFC with 5-second interval), and HC (homecage) groups. **(C)** Representative image of RBs in the BLA. Mean (± standard error) of double-labeled (c-Fos-positive and RB-positive) cells (left, striped), c-Fos-positive cells (middle, grey), and RB-positive cells (right, white) in the **(D)** prelimbic cortex **(PL), (E)** ventral CA1 (vCA1), **(F)** perirhinal cortex (PER), and **(G)** lateral entorhinal **cortex (LEC).** * Indicates p < 0.050. Generalized Linear Models followed by LSD test. Dots show sample distribution.

### 3.5 CFC-5s learning induced higher activation of the PL

The c-Fos expression was consistent and replicated those observed in Experiment 2. There was a significantly higher c-Fos expression in the PL and vCA1 induced by CFC-5s training compared to CFC and HC groups and induced by CFC training compared to the HC group. Both fear conditioning had a significantly higher c-Fos expression in the PER, LEC, AC, IL, and RE than in the HC group, as previously observed in Experiment 2 (Figures 4D-G and Figures 5A-D). GZLM showed an effect of the group for c-Fos-positive cells in the PL (W = 50.583; df = 2; p = 0.001), vCA1 (W = 16.655; df = 2; p = 0.001), PER (W = 32.419; df = 2; p = 0.001), LEC (W = 61.207; df = 2; p = 0.001), AC (W = 26.685; df = 2; p = 0.001), IL (W = 18.178; df = 2; p = 0.001), RE (W = 21.569; df = 2; p = 0.001), and vSUB (W = 10.017; df = 2; p = 0.007), but not in the VS (W = 0.373; df = 2; p = 0.830). LSD test showed that the CFC-5s group had significantly higher c-Fos expression than those in the PL (p = 0.048; β = 0.614) and vCA1 (p = 0.038; β = 0.843). Both CFC and CFC-5s groups had significantly higher c-Fos expression than the HC group in the PL (CFC p = 0.001; β = 1.324; CFC-5s p = 0.001; β = 1.938), vCA1 (CFC p = 0.002; β = 0.915; CFC-5s p = 0.001; β = 1.757), PER (CFC p = 0.001 β = 1.552; CFC-5s p = 0.001; β = 1.616), LEC (CFC p = 0.001; β = 1.816; CFC-5s p = 0.001; β = 1.458), AC (CFC p = 0.001; β = 1.424; CFC-5s p = 0.001; β = 1.139), IL (CFC p = 0.001; β = 1.607; CFC-5s p = 0.002; β = 1.110), RE (CFC p = 0.001 β = 1.358; CFC-5s p = 0.001; β = 1.244), and vSUB (CFC p = 0.002; β = 1.363; CFC-5s p = 0.021; β = 1.037).

**Figure 5.**
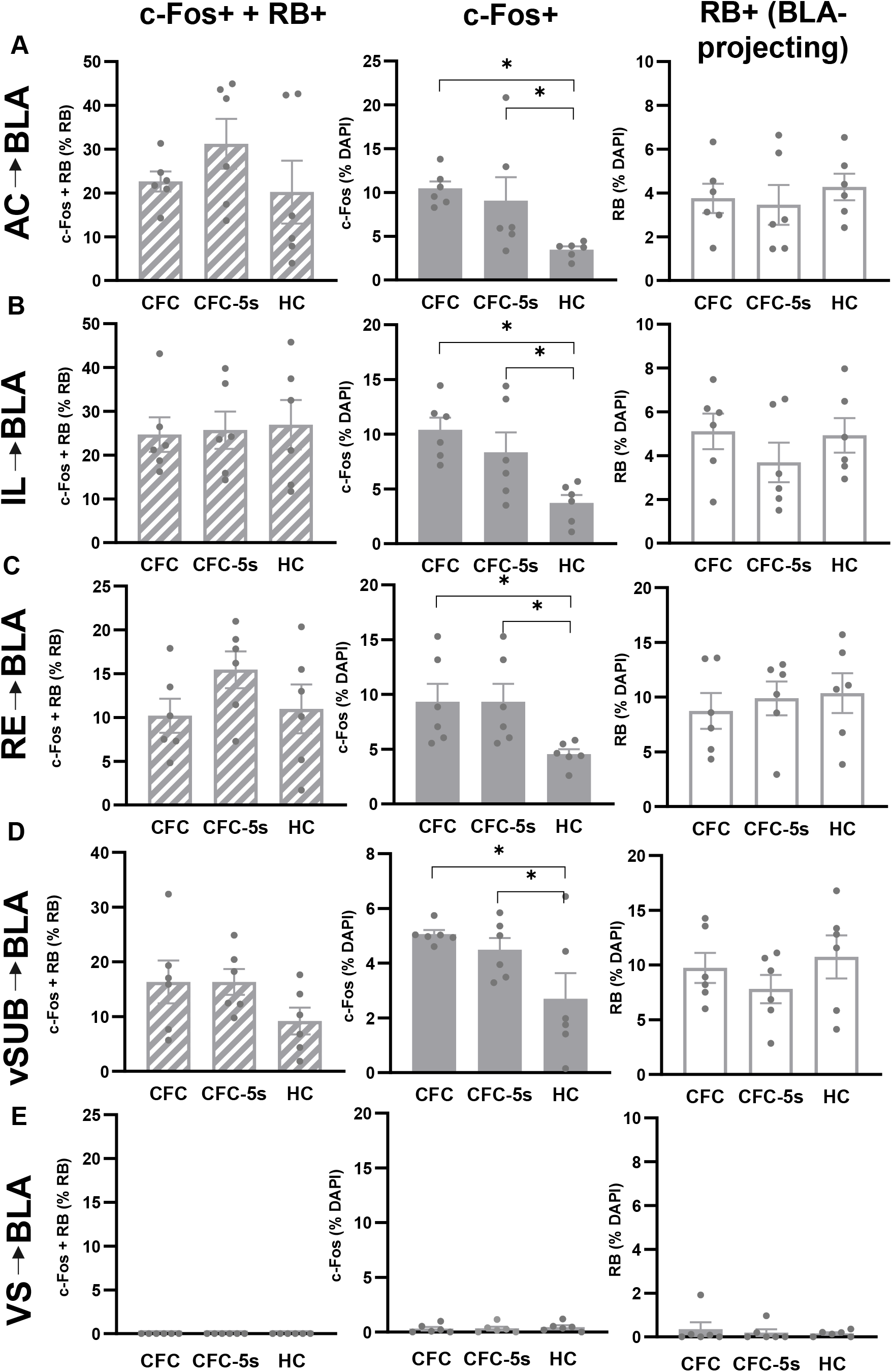
CFC and CFC-5s learning activated cells in the AC, IL, RE, and vSUB, replicating the previous results. Mean (± standard error) of double-labeled (c-Fos-positive and RB-positive) cells (left, striped), c-Fos-positive cells (middle, grey), and RB-positive cells (right, white) in the **(A)** cingulate cortex (AC), **(B)** infralimbic cortex (IL), **(C)** nucleus reuniens of the thalamus (RE), **(D)** ventral subiculum (vSUB), and **(E)** ventral striatum (VS). * Indicates p < 0.050. Generalized Linear Models followed by LSD test. Dots show sample distribution.

All groups had similar freezing times during the training session (Figure 4A) and an equal average of cells (DAPI-positive cells) and PL-projecting cells in all brain regions. GZLM did not show an effect of the group in the mean freezing time in the training session (W = 0.272; df = 1; p = 0.602) nor in DAPI-positive cells in the PL (W = 0.044; df = 2; p = 0.978), vCA1 (W = 4.871; df = 2; p = 0.088), PER (W = 1.425; df = 2; p = 0.491), LEC (W = 0.455; df = 2; p = 0.796), AC (W = 0.184; df = 2; p = 0.912), IL (W = 0.227; df = 2; p = 0.893), RE (W = 1.599; df = 2; p = 0.449), vSUB (W = 2.080; df = 2; p = 0.353), and VS (W = 2.936; df = 2; p = 0.230). GZLM also did not show an effect of group in RB-positive cells in the PL (W = 0.675; df = 2; p = 0.714), vCA1 (W = 0.429; df = 2; p = 0.807), PER (W = 1.251; df = 2; p = 0.535), LEC (W = 0.089; df = 2; p = 0.957), AC (W = 0.721; df = 2; p = 0.697), IL (W = 2.005; df = 2; p = 0.367), RE (W = 0.603; df = 2; p = 0.740), vSUB (W = 2.076; df = 2; p = 0.354), and VS (W = 1.080; df = 2; p = 0.583). Figures 4B and 4C illustrate the diffusion pattern of the RBs in the PL. Grey areas represented the locations of the micro-infusions.

Overall, these results indicate that activation of PL projections to the BLA and increased activation of the PL and vCA1 underlie CFC-5s learning, whereas activation of vCA1 projections to the BLA, and increased activation of the PER, LEC, AC, IL, RE, and vSUB underlie CFC learning, despite of the time interval. These results indicate that differential labeling of BLA-projecting neurons among the groups could not explain the results. In addition, BLA-projecting neurons in the VS were also undetected, suggesting specific BLA targeting and labeling of BLA-projecting neurons.

#### 4. Discussion

We investigated the activation of direct projections to the PL or the BLA induced by CFC and CFC-5s learning, fear associations of stimuli overlapped and separated in time, by quantifying the c-Fos expression following CFC or CFC-5s training in neurons from 9 brain regions projecting to the PL or BLA, identified with the retrograde trace RBs. Results showed that the CFC-5s training activated the vCA1→PL, PER→PL, and PL→BLA monosynaptic connections. Both the CFC and CFC-5s training activated the BLA→PL, vCA1→BLA, and LEC→PL monosynaptic connections. Moreover, the CFC-5s training activated the PL and vCA1, and both the CFC-5s and CFC training the BLA, PER, LEC, AC, IL, RE, and vSUB. The temporal association engaged more PL neuroanatomical connections, and both fear associations BLA neuroanatomical connections.

PL afferences could converge the CS to be represented during the time interval, and PL efferences could send a CS representation to be associated with the US afterward. In this scenario, we hypothesize that the PER and the vCA1 would convey the contextual CS to the PL, and the PL would convey the contextual CS to the BLA to be associated with the US in the CFC-5s learning.

We have previously shown that the functional disconnection of the vCA1 and PL by pre-training pharmacological asymmetric inactivation impaired the CFC-5s but not CFC learning (Santos et al. 2020). Here, we expanded these results by showing that this functional connection involves the activation of the vCA1→PL projection. Thus, observational and confirmatory evidence suggests a functional and neuroanatomical connection between the vCA1 and PL in CFC-5s learning. Our results also corroborated previous findings showing no increase in c-Fos expression in vCA1 PL-projecting neurons after the CFC training compared to a control group for contextual learning (Dixsaut and Gräff 2022).

Given that the DH is required for contextual learning (Fanselow 2010) but has undetected or sparse direct projections to the PL (Hoover and Vertes 2007; Kim and Cho 2017; Ye et al. 2017), the VH could convey contextual information to the PL to be maintained during the time interval in the CFC-5s learning. In line, the VH is also required for contextual learning (Rudy and Matus-Amat 2005) and CFC (Bast et al. 2001; Zhang et al. 2001; Kim and Cho 2020). Both contextual learning and CFC increased Arc (activity-regulated cytoskeleton-associated protein) in the vCA1 (Czerniawski et al. 2011; Hudgins and Otto 2019). The dCA1 and vCA1 have engram cells, that is, cells activated by the CFC learning and reactivated by memory retrieval (Tayler et al. 2013; Trouche et al. 2013; Roy et al. 2022). CA1 neurons activated by the contextual CS were preferentially reactivated following memory retrieval, and their artificial reactivation facilitated and their inhibition impaired the contextual fear memory, suggesting that the CA1 encodes the context that is fear-associated in CFC (Deng et al. 2013; Nakazawa et al. 2016; Ghandour et al. 2019). Due to its direct connection to the PL, the CFC-5s may engage more the vCA1 than the CFC, as shown by increased c-Fos expression.

The neuroanatomical connections between the vCA1 and PL are also required for the retrieval of contextual fear memory (Hallock et al. 2020), fear renewal (Orsini et al. 2011; Jin and Maren 2015; Wang et al. 2016; Vasquez et al. 2019), and SWM (Spellman et al. 2015). SWM is also accompanied by synchronized theta activity between the VH and mPFC (Jones and Wilson 2005; Siapas et al. 2005; Hyman et al. 2010; O’Neill et al. 2013; Spellman et al. 2015; Hallock et al. 2016), and their functional disconnection impaired learning, especially with more extended delay periods (Floresco et al. 1997; Wang and Cai 2006; Churchwell and Kesner 2011). On the other hand, silencing inputs from vCA1 to the PL during the training session did not impair trace fear conditioning but decreased the retrieval of background CFC in the first minutes of the test session, which was overcome with additional time or further training (Twining et al. 2020). These findings suggest that combined temporal and spatial factors, such as in CFC-5s and SWM, but not temporal or spatial factors alone require the vCA1→PL connection. The vCA1 may intentionally convey spatial cues or incidentally transfer past contextual states of the animal to the PL, which would only become essential when there were no other salient and predictive stimuli available to be associated, as in the CFC-5s. Alternatively, the involvement of the vCA1-PL functional connection in CFC-5s could reflect a function in anxiety-related states (Schoenfeld et al. 2014; Ciocchi et al. 2015; Padilla-Coreano et al. 2016).

In addition to the VH, the PER→PL projection could indirectly connect the DH with the PL. The PER integrates inputs from all sensory modalities and projects directly to the CA1 and subiculum, which projects back to them (Burwell and Amaral 1998). The PER also indirectly projects to the CA1 and subiculum and the other hippocampal subdivisions via the LEC (Doan et al. 2019), which conveys non-spatial stimuli, such as visual, olfactory, and object information in the so-called “what pathway” (Furtak et al. 2007). The DH also has time cells, neurons that fire during specific periods in temporally structured episodes observed during tasks involving a sequence of events (MacDonald et al. 2011). In addition, dCA1 neurons showed CS-related responses in the training and test session of trace conditioning, which was positively correlated with the exhibition of the CR (Zhang et al. 2019; Ahmed et al. 2020; Mount et al. 2021). Therefore, although the VH can support contextual learning and CFC, temporal associations may involve other functions specifically related to the DH and indirect connections with the PL, such as via the PER. A functional connection between the DH and the PL accompanied CFC-5s learning. Pre-training PL inactivation impaired CFC-5s learning and reduced CFC-5s-induced phosphorylation of CREB in the DH (Santos et al. 2023). The PER was a hub, a brain region with increased importance, in the functional network of CFC learned in the absence of the DH, that is, with a dorsal hippocampal lesion, suggesting that the PER may replace the DH exercising a similar function (Coelho et al. 2018).

The PER can organize individual elements in configural representations, such as contexts. The PER is required for CFC, in line with the increased c-Fos expression in the PER by both CFC and CFC-5s learning (Bucci et al. 2000; Lindquist et al. 2004; Kholodar-Smith et al. 2008a; Kholodar-Smith et al. 2008b). The PER is also necessary for AFC using ultrasonic vocalization of rats (USVs) or discontinuous tones, in which tone segments need to be unitized (Lindquist et al. 2004; Furtak et al. 2007; Kholodar-Smith et al. 2008a), contextual discrimination (Bucci et al. 2002) and object recognition (Winters and Bussey 2005; Barker et al. 2007; Bartko et al. 2007; Hernandez et al. 2017), both which require conjunctive representations of the context and objects, respectively, to solve the ambiguity, given that recognizing or discriminating objects or contexts could not be done based on their elements due to overlapping features. In the CFC-5s learning, the PER could support the formation of a configural representation of the context and convey it to the PL via its direct projections. Moreover, the PER has endogenous persistent-firing neurons (Navaroli et al. 2012) and is required for trace fear conditioning (Kholodar-Smith et al. 2008b; Suter et al. 2013), which showed increased PER-responsive neurons (Suter et al. 2019). Therefore, the PER could have a role in stimulus unitization, assembling the context based on CFC, object recognition, contextual discrimination, and fear conditioning using USV or discontinuous tones and in transient memories based on its endogenous persistent activity.

Functional disconnection of the PER and mPFC was required in detecting and discriminating object-in-place associations, which involves flexibly updating which object is rewarded depending on the animal’s location (Hernandez et al. 2017). In the CFC-5s learning, activation of the PER→PL connection may underlie the detection of a novel stimulus-context association, updating that the context previously neutral is after the interval threatening. In both tasks, there was a change in the stimuli associations, which was required to adapt the animal’s response. In line, the PL was needed to successfully encode context-safety contingencies in a context fear discrimination task (Corches et al. 2019) and contributed to learning compound stimulus-reward associations in a set-shifting task (Mukherjee and Caroni 2018), suggesting that it could encode functional values as context-safety and context-danger relationships in the CFC-5s learning. Activation of the PER→PL connection may underlie the detection of association changes to adapt the behavioral response. Thus, the PER→PL could convey the context from the DH; from the PER itself, based on its unitizing of stimuli; or update the context-stimulus association. The PER→PL and vCA1→PL could be redundant or complementary circuits. The functional disconnection of the vCA1-PL impaired CFC-5s learning, suggesting that the PER-PL connection would not be sufficient to replace it or that it is not independent of the vCA1-PL functional connection. Future gain- or loss-of-function studies investigating the functionality of PER-PL connections would address if they were also necessary for CFC-5s learning.

In turn, activating vCA1→BLA projections accompanied the CFC and CFC-5s learning. Thus, depending on their projection targets, vCA1 neurons are engaged in contextual fear associations separated or overlapped in time. PL-projecting vCA1 neurons were recruited in temporal associations, and BLA-projecting vCA1 neurons in both fear associations. In CFC, the vCA1 conveys the contextual CS to be associated with the US in the BLA. CFC induced synaptic strengthening between vCA1 neurons projecting to the BLA. BLA neurons received inputs from vCA1 neurons activated by the context, and the opto-inhibition of these context-responsive vCA1 neurons impaired the CFC and reduced the synaptic strengthening in the vCA1→BLA connection. Moreover, activation of the vCA1→BLA connection paired with the US promoted CFC, whereas its chemogenetic inhibition impaired CFC learning (Kim and Cho 2020). The opto-inhibition of BLA-projecting neurons in the vCA1 or the opto-stimulation of vCA1 terminals in the BLA, which disrupts its regular activity, also impaired CFC learning (Xu et al. 2016; Jimenez et al. 2018).

The vCA1 has neurons that simultaneously project to both PL and BLA with axon collaterals, facilitating synchronized activity (Ishikawa and Nakamura 2006). These neurons preferentially engage in contextual learning (Kim and Cho 2017) and fear renewal (Jin and Maren 2015). CFC-5s learning activated the vCA1→PL, vCA1→BLA, and PL→BLA connections. vCA1 double-projecting neurons could convey the contextual CS directly to the PL and BLA and indirectly through the PL→BLA connection. Coordinated activity between the vCA1, PL, and BLA could form a recurrent circuit that provides reverberant activity related to the context (Wang 2001). It remains to be determined whether single- or double-projecting neurons were activated in the vCA1 following CFC-5s learning and whether PL neurons receiving inputs from vCA1 and projecting to the BLA are activated.

The PL was the only brain region among the 9 investigated with higher activation of BLA-projecting neurons following CFC-5s learning. Considering that BLA is the site of CS-US convergence (Barot et al. 2009), this finding suggests that the PL may convey the contextual CS associated with the US in the BLA. In line, a functional connection between the PL and the BLA accompanied CFC-5s learning (Santos et al. 2023), and the PL→BLA connection was required or accompanied trace fear conditioning, but not CFC (Song et al. 2005; Kirry et al. 2020; Kitamura et al. 2017). Olfactory or AFC also induced post-training changes in the PL-BLA functional connection required for the expression of the CR (Garcia et al. 1999; Vouimba et al. 2011; Arruda-Carvalho and Clem 2014; Karalis et al. 2016). The chemogenetic silencing of PL inputs to the BLA impaired trace fear conditioning with 2, but not 6, CS-US pairings (Kirry et al. 2020). This finding suggests that when the association is not formed, the BLA can rely on stimuli represented in PL, and cortical input influences CS selection. Once learned the contingency, activation of PL projections to the BLA may be unnecessary. This can be specialty true when no other salient stimulus exists, such as in the CFC-5s learning, which also has 1 CS-US pairing.

Both CFC-5s and CFC learning activated the BLA→PL and LEC→PL connections, which could underly a system consolidation for remote memories. During the CFC training, silencing of the BLA or medial entorhinal cortex (MEC) terminals in the PFC impaired the remote, but not recent, memory retrieval (Kitamura et al. 2017). CFC learning also increased c-Fos expression in BLA and EC neurons projecting to the PL compared to contextual learning, and the pre-training chemogenetic inhibition of the BLA→PL and EC→PL connections impaired the remote, but not recent retrieval of the contextual fear memory (Dixsaut and Gräff 2022). These results suggest that BLA and EC connections to PL during learning are functional for later memory retrieval.

CFC and CFC-5s learning also induced higher c-Fos expression in the BLA and EC, in agreement with their participation in CFC learning. Pre-training or post-training lesions, inactivation, or NMDA receptor antagonism in the BLA impaired CFC (Maren et al. 1996; Goosens and Maren 2003; Matus-Amat et al. 2007). The BLA also has engram cells activated by CFC learning and memory retrieval (Reijmers et al. 2007; Tayler et al. 2013; Nonaka et al. 2014; Kitamura et al. 2017; Roy et al. 2022). In turn, pre-training lesion (Maren and Fanselow 1997; Majchrzak et al. 2006), pre-training NMDA receptor antagonism (Schenberg et al. 2005), or post-training inactivation (Baldi et al. 2013) of the EC impaired CFC. CFC training also increased the EC’s ERK (extracellular signal-regulated kinase) (Hebert and Dash 2004).

Finally, we also observed higher RE, AC, and IL activation induced by CFC and CFC-5s training. The RE is required for recent fear generalization (Xu and Sudhof 2013; Wu and Chang 2022), remote memory retrieval of CFC (Wheeler et al. 2013; Vetere et al. 2017), and remote retrieval of spatial memories (Loureiro et al. 2012; Klein et al. 2019). The opto-stimulation of encoding ensembles in the RE increased the CR, suggesting that it may support CFC memory formation (Roy et al. 2022). Thus, increased c-Fos expression in the RE may underly later remote memory retrieval. Regarding temporal learning, pre-training inactivation or high-frequency stimulation of the RE during the interval impaired trace fear conditioning (Eleore et al. 2011; Lin et al. 2020; Wu and Chang 2022; Jhuang and Chang 2023). In turn, the AC is required for the recent and remote consolidation of the CFC. Post-training changes, such as NR2A transmission, protein synthesis, and transcriptional and structural remodeling in the AC, are necessary for the consolidation of contextual fear memories (Tang et al. 2005; Zhao et al. 2005; Vetere et al. 2011; Einarsson and Nader 2012; Bero et al. 2014). The IL is classically required for fear extinction (Do-Monte et al. 2015), but contextual learning and CFC also increase zif268 (zinc finger protein 268) in the IL (Asok et al. 2013; Chakraborty et al. 2016).

C-Fos expression was consistent across the experiments in the 8 brain regions that were double investigated (AC, IL, LEC, PER, RE, vCA1, vSUB, VS). The same activation pattern among the groups observed in Experiment 2 was replicated in Experiment 3. The CFC and CFC-5s groups had higher c-Fos expression than the HC in the AC, IL, LEC, PER, RE, vSUB, and vCA1; the CFC-5s also than the CFC in the vCA1, and all groups had similar c-Fos expression in the VS, in both Experiment 2 and 3. Double-labeled cells could occur by chance in RB-positive cells unrelated to their functionality. The higher the c-Fos- or RB-positive cells, the higher the probability of them co-occurring randomly. However, c-Fos expression and double-labeled cells did not appear to correspond in our study. For instance, the CFC-5s group had higher c-Fos expression in the vCA1 than the CFC and HC groups in Experiments 2 and 3, but only in Experiment 3, they had higher double-labeled cells than the CFC and HC groups. Both CFC-5s and CFC groups had higher c-Fos expression in the PER than the HC, but only the CFC-5s group had higher double-labeled cells than the HC group. The CFC-5s and CFC groups also had higher c-Fos expression than the HC group in all the brain regions except the VS. However, in any group, we did not observe higher double-labeled cells than the HC in the AC, the IL, the RE, or the vSUB.

To help ensure that the infusion of the retrograde tracer was restricted to target brain regions of interest and minimal to fibers of passage, we verified the labeling of RB-positive cells in the VS, a brain region that does not receive projections from the PL or the BLA. Results showed that RB-positive cells in the VS were extremely low to undetected, suggesting that the labeling observed was majorly specific to PL- and BLA-projecting cells and that contamination of fibers of passage in the PL or BLA had minimal impact and did not represent a significant source of artifact, being similar among the groups.

Present results highlighted the relevance of the vCA1→PL, the PER→PL, the PL→BLA connections, and the vCA1 and PL in temporal associations. Associations separated in time involved additional neuroanatomical connections than associations overlapped in time. Multiple pathways may participate due to the absence of salient stimulus, uncertainty, and engagement of a transient memory system, suggesting a more complex association. Based on the functions of brain regions and projections, results helped better understand the learning mechanisms of temporal associations, adding the relevance of the PER→PL connection for the first time.

## Acknowledgments

This work was supported by São Paulo Research Foundation (FAPESP) grant number #2016/25755-2, #2016/13027-2, and #2019/04844-5; Conselho Nacional de Desenvolvimento Científico e Tecnológico (CNPq); Coordenação de Aperfeiçoamento de Pessoal de Nível Superior (CAPES) and Associação de Fundo de Incentivo à Pesquisa (AFIP).

